# Molecular basis of mitotic decline during human aging

**DOI:** 10.1101/261008

**Authors:** Joana Catarina Macedo, Sara Vaz, Bjorn Bakker, Rui Ribeiro, Petra Bakker, Jose Miguel Escandell, Miguel Godinho Ferreira, René Medema, Floris Foijer, Elsa Logarinho

## Abstract

Aneuploidy, an abnormal chromosome number, has been linked to aging and age-associated diseases, but the underlying molecular mechanisms remain unknown. Supported by direct live-cell imaging of young, middle-aged and old-aged primary human dermal fibroblasts, we found that aneuploidy increases with aging due to general dysfunction of the mitotic machinery. Increased chromosome segregation defects in elderly mitotic cells correlated with an early senescence-associated secretory phenotype (SASP) and repression of Forkhead box M1 (FoxM1), the transcription factor that drives expression of most G2/M genes. By restoring FoxM1 levels in elderly and Hutchison-Gilford Progeria Syndrome fibroblasts we prevented aneuploidy and, importantly, ameliorated cellular phenotypes associated with aging. Moreover, senescent fibroblasts isolated from elderly donors’ cultures were mostly aneuploid, suggesting that aneuploidy is a key player in the progression into full senescence phenotypes. Based on this feedback loop between cellular aging and aneuploidy, we propose modulation of mitotic efficiency through FoxM1 as a potential strategy against aging and progeria syndromes.

---

Aging has been linked to an increase in aneuploidy for the past several decades^1^. This association has been well documented for oocytes and is considered to be the main cause of miscarriage and birth defects in humans^2^. However, aneuploidy can also arise in somatic cells, and a number of studies have reported age-dependent increases in aneuploidy^3–7^. These studies have shown that aging is positively correlated with the incidence of chromosome mis-segregation, raising the question whether there is a general dysfunction of the mitotic apparatus in aged cells^8^. Transcriptome analyses of a panel of fibroblast and lymphocyte cultures from young and old individuals revealed changes in the expression of genes controlling the mitotic machinery^9,10^. However, the use of mixed cell populations in different stages of the cell cycle and the lower mitotic indexes of elderly cell cultures has limited these studies from clearly demonstrating whether mitotic genes are repressed intrinsically in old dividing cells. Moreover, analysis of the mitotic process in aging models and diseases has been scarce and a comprehensive analysis of cell division efficiency in naturally aged cells is largely missing.

More recently, aneuploidy has been also linked to aging. Studies of aneuploidy-prone mouse models exhibiting increased rates of chromosome mis-segregation uncovered a unanticipated nexus with the rate of aging and the development of age-related pathologies^11–14^. Mutant mice with low levels of the spindle assembly checkpoint protein BubR1 were found to develop progressive aneuploidy along with a variety of progeroid features, including short lifespan, growth retardation, sarcopenia, cataracts, loss of subcutaneous fat and impaired wound healing ^11^. In addition, systematic analyses of disomic yeast, trisomic mouse and human cells, all cells with an extra chromosome, have elucidated the impact of aneuploidy in cellular fitness^15^. The cumulative effect of copy number changes of many genes induces the so called pan-aneuploidy phenotypes^16^, namely proliferation defects^17,18^, gene signature of environmental stress response^19,20^, multiple forms of genomic instability^21–23^, and proteotoxicity^24–26^, which interestingly, are cellular hallmarks of aging^27^.

Together, these observations suggest that there is a positive correlation between aging and aneuploidy but evidence for mitotic decline in elderly dividing cells and for aneuploidy-driven permanent loss of proliferation capacity (full senescence) is limited. Here, we used live cell time-lapse imaging to investigate the mitotic behavior of human dermal fibroblasts collected from healthy Caucasian males with ages ranging from neonatal to octogenarian and cultured under low passage number. We show mitotic duration to increase with advancing age alongside with a higher rate of mitotic defects. We demonstrate this mitotic decline to arise from a transcriptional shutdown of mitotic genes in pre-senescent dividing cells exhibiting senescence-associated secretory phenotype (SASP). We show short-term induction of FoxM1 transcriptional activity to improve mitotic fitness and ameliorate senescence phenotypes in elderly and progeroid cells. Finally, we identify aneuploidy as a major hallmark of full senescence in naturally aged cells thus pointing up for the potential of mitotic fitness enhancement as an anti-aging strategy.

## Results

### Aneuploidy increases during natural aging

In our study we used human dermal fibroblasts (HDFs) collected from healthy Caucasian males with ages ranging from neonatal to octogenarian (Supplementary Fig.1a), to test whether aneuploidy increases during natural aging and if this is a consequence of a general dysfunction of the mitotic machinery in aged cells. Since inter-individual differences exist in the rate at which a person ages (biological age), we included two donors of each age in a total of five chronological ages to increase the robustness of any correlation found with chronological aging. In addition, we used dermal fibroblasts from an 8 years-old child with the Hutchison-Gilford progeria syndrome (HGPS) as a model of premature cellular aging^28^. Considering that during a normal post-natal lifespan cells *in vivo* will hardly reach the exhaustion number of replications observed in culture^29^, and to limit any artifacts and clonal selection of *in vitro* culturing, only early cell culture passages way below replicative lifespan exhaustion were used (Population doubling level (PDL)<24). Chronic accumulation of macromolecular damage during natural aging induces a cellular stress response known as senescence^30^, and accumulation of senescent cells has been widely used to reflect biological age and linked to pathological aging^31,32^. Accordingly, we found significantly higher levels of senescence-associated (SA) biomarkers^33^, measured under strict quantitative parameters by microscopy analysis, in the proliferating fibroblast cultures from older individuals and the HGPS patient (Supplementary Fig.1b-g), thus validating their suitability as models of advanced and premature aging, respectively.

To determine if aneuploidy increases with aging, we performed fluorescence *in situ* hybridization (FISH) analysis for three chromosome pairs. We found significantly higher aneusomy indexes in asynchronous cell populations of middle-aged, old-aged and progeria samples (Fig. 1a,b). However, because aneuploid cells are most likely outcompeted in culture by diploid cells^17^, thereby diluting the aneuploidy index, we additionally measured aneuploidy in a post-mitotic cell subpopulation generated during a 24h treatment with the cytokinesis inhibitor cytochalasin D (Fig. 1a,c). Using this approach, we observed even higher aneuploidy in the middle-aged, old-aged and progeria cultures. Thus, the mild aneuploidy levels found in elder cultures support the idea of an age-associated loss of mitotic fidelity.

**Figure 1.**
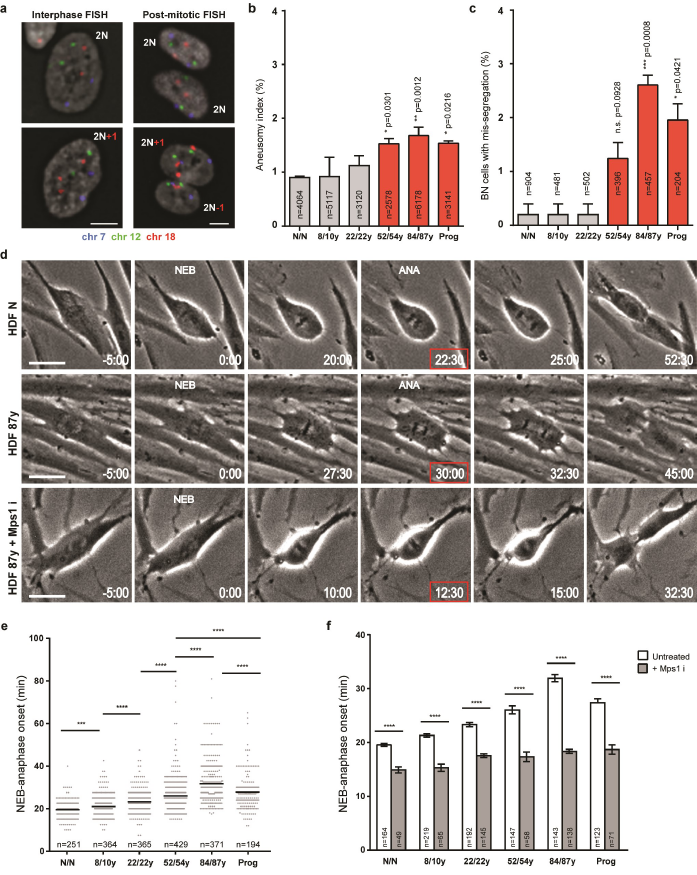
Aneuploidy and mitotic duration increase during normative aging. **(a)** FISH in interphase and post-mitotic cells with chromosome-specific centromeric probes as indicated. Representative images of euploid/2N and aneuploid/2N±l cells are shown. Scale bars, 5 µm. **(b)** Aneusomy index in interphase cells from different age donors. **(c)** Percentage of post-mitotic cells with chromosome mis-segregation. **(d)** Frame series of time-lapse phase-contrast movies of mitotic neonatal and elderly cellsMps1 inhibitor. Mitotic duration was measured from nuclear envelope breakdown (NEB) to anaphase onset (ANA). Time, min:sec. Scale bar, 20 µm. **(e)** Mitotic duration of individual fibroblasts from different donors. **(f)** Mitotic duration under standard conditions as in (e) (white bars) and following treatment with Mps1 inhibitor (gray bars). Values are mean ± s.d. from at least three independent experiments, using two biological samples of similar age (except for progeria). Sample size (n) is indicated in each graph. n.s. p>0.05, *p≤0.05, **p≤0.01, ***p≤0.001 and ****p≤0.0001 in comparison to neonatal (N/N) by two-tailed χ^2^ (b,c) and Mann-Whitney (e,f) statistical tests.

### Elder cells divide slower with increased rate of mitotic defects

To gain insight into how old cells divide, we followed individual mitotic cells by long-term phase-contrast time-lapse imaging. Interestingly, we found the interval between nuclear envelope breakdown (NEB) and anaphase onset to increase steadily with advancing age (Fig. 1d-e). To exclude any effects due to genetic heterogeneity between the Caucasian donors and to discrepancies in culture population doubling levels, we used mouse adult fibroblasts recurrently sampled from female mice over a period of 2 years. In this model, culture conditions were highly controlled, thereby providing a solid “ex vivo” model of chronological aging. Again, both mitotic duration (Supplementary Fig.2a-c) and senescence-associated biomarkers (Supplementary Fig.2d-e) progressively increased with aging.

We then asked what could be leading to the age-associated mitotic delay. Bypass of short telomere-triggered senescence by disruption of tumor-suppressive pathways, shown to elicit telomere fusion-driven prolonged mitosis^34^, was ruled out as potential cause since we found no evidence for chromosome fusions in metaphase spreads of the primary dermal fibroblasts used in our study (Supplementary Fig.3a). Activation of the spindle assembly checkpoint (SAC) by defective kinetochore-microtubule attachments and/or reduced efficiency of the ubiquitin-proteasome system induced by proteotoxic stress could alternatively explain the delay between NEB and anaphase onset in older cells. We found the mitotic delay to depend on SAC activity as treatment with a small molecule inhibitor of the Mps1 kinase rescued mitotic duration to similar levels in all cell cultures (Fig. 1d,f). Enhancement of proteasome activity with a small molecule inhibitor of the Usp14 deubiquitinase did not change mitotic duration considerably (Supplementary Fig.3b). As a correlation between cell size, mitotic duration and SAC strength was recently described^35^, we further tested whether aging-associated mitotic delay was due to increased cell size. We found no significant correlation between mitotic duration and cell size in fibroblast cultures (Supplementary Fig.3c-f). In addition, when treated with the kinesin-5 inhibitor (STLC) to induce chronic activation of the SAC, elderly cells arrested in mitosis as long as young cells before they slipped out, suggesting SAC strength is similar (Supplementary Fig.3g).

To investigate chromosome and/or spindle defects contributing to increased mitotic duration in older cells, we performed high-resolution spinning-disk confocal microscopy in cells expressing H2B-GFP and -Tubulin-mCherry. We found several mitotic defects to be significantly increased in middle-aged and old-aged samples (Fig. 2a-d), namely chromosome congression delay (Fig. 2b), anaphase lagging chromosomes (Fig. 2c) and spindle mispositioning in relation to the growth surface (Fig. 2d). These phenotypes were quantified by scoring the percentage of cells exhibiting severe vs. mild chromosome mis-segregation following treatment with the Mps1 inhibitor (Fig. 2e; Supplementary Fig.4a); micronuclei^36^ (Fig. 2f); spindle mispositioning (Fig. 2g, Supplementary Fig.4b-d); and cytokinesis failure (Fig. 2h). Overall, the data indicated that alongside increased aneuploidy, aging triggers abnormalities at several mitotic stages.

**Figure 2.**
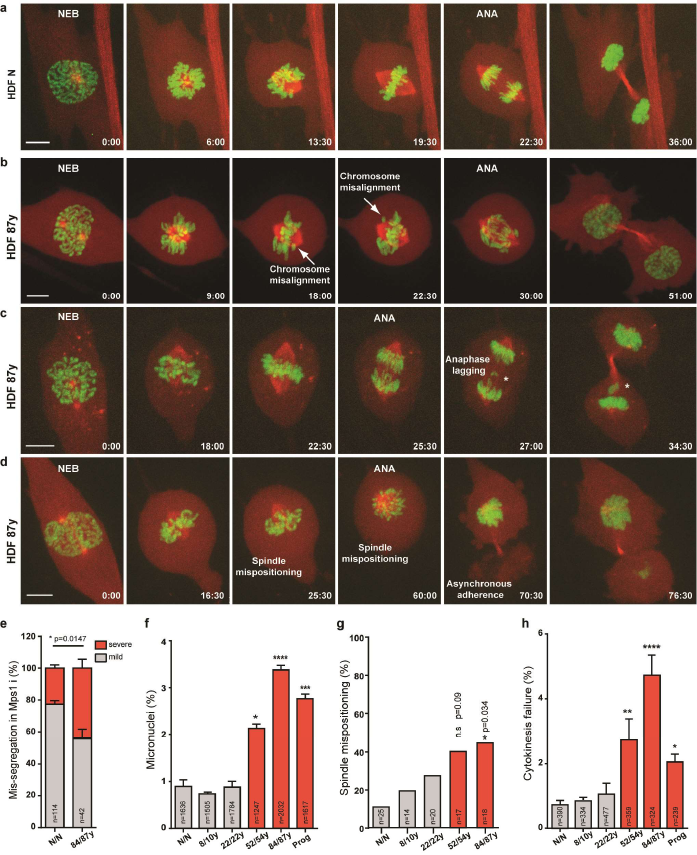
Mitotic defects associated with advanced age. **(a-d)** Frame series of spinning-disk confocal movies of neonatal (HDF N) and elderly (HDF 87y) dividing cells expressing H2B-GFP--Tubulin-mCherry. **(a)** Neonatal cell division. **(b-d)** Elderly cell with chromosome alignment delay **(b)**, with anaphase lagging chromosome and micronucleus formation (*) **(c)**, or with spindle mispositioning in relation to the growth surface and asynchronous adherence of the daughter cells **(d)** (Supplementary Video 1). Time, min:sec. Scale bar, 5 µm. **(e)** Percentage of cells with mild and severe chromosome mis-segregation following treatment with Mps1 inhibitor. **(f)** Percentage of cells with micronuclei. **(g)** Fraction of cells with spindle mispositioning. **(h)** Rate of cytokinesis failure. Values are mean ± s.d. from at least two independent experiments, using two biological samples of similar age (except for progeria). Sample size (n) is indicated in each graph. *p≤0.05, **p≤0.01, ***p≤0.001 and ****p≤0.0001 in comparison to N/N by two-tailed χ^2^ (e-h) and Mann-Whitney (j) statistical tests.

### Elderly mitotic fibroblasts exhibit SASP and transcriptional shutdown of mitotic genes

To identify the molecular mechanisms behind this complex aging-associated mitotic phenotype, we performed RNA sequencing (RNA-seq) gene expression profiling of cells captured in mitosis accordingly to the experimental layout shown in Fig. 3a. This procedure yielded purified cell populations with mitotic indices >95% in both neonatal and 87y donor cultures (Fig. 3b). RNAseq revealed that the abundance of 3,319 gene transcripts was significantly altered in octogenarian mitotic fibroblasts compared to neonatal mitotic fibroblasts (Fig. 3c). Principal component analysis (PCA) showed that the biological replicates were consistent (Fig. 3d). In agreement with the elderly cell defects in mitosis, the top 10 most altered Gene Ontology (GO) terms included 7 cell cycle- and mitosis-related gene ontologies (Fig. 3e). 105 ‘mitotic cell cycle’ genes were significantly altered in octogenarian mitotic fibroblasts (p-value<0.05; Supplementary Table 1), representing a 1.7-fold enrichment of alterations in this gene set (*p*-value=8.4E-9) (Fig. 3g). Moreover, we found significantly altered expression of 16 out of 29 cytokines, metalloproteinases (MMPs) and other factors previously associated with the senescence-associated secretory phenotype (SASP)^37^, in the elderly vs. neonatal mitotic cells (Fig. 3f). Thus, an unforeseen SASP phenotype evolves in elderly dividing cells, alongside a global transcriptional shutdown of mitotic genes, likely accounting for aging-associated mild aneuploidy levels.

**Figure 3.**
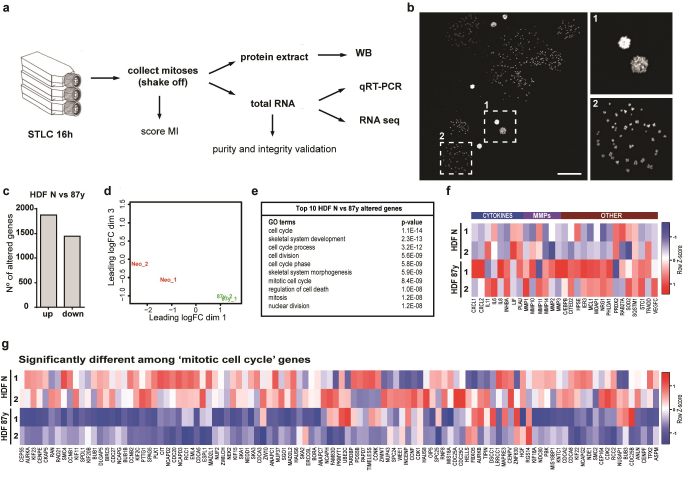
Elderly mitotic fibroblasts exhibit a downregulation of cell cycle genes. **(a)** Experimental layout. Fibroblast cultures from different age donors were treated with kinesin-5 inhibitor (STLC) to enrich for mitotic index (MI). Mitotic (detached) cells were collected and inspected for MI>95%. Protein extracts and total RNA were prepared for gene expression analyses. (b) Representative image of MI>95%. Scale bar, 100 µm. (c) Number of altered genes of HDF 87y, comparing to HDF N. (d) Multidimensional scaling plot of distances between gene expression profiles based on sample RNA-sequencing. (e) Top ten altered GO terms organized by p-values using the DAVID Functional Annotation tool. (f) Heatmap of SASP gene (cytokines, metalloproteinases (MMPs) and other factors) expression in HDF N and HDF 87y. Genes are represented in columns and biological replicates are represented in rows. In the Z-score column, color intensities represent higher (red) to lower (blue) expression. (g) Heatmap of genes within the ‘mitotic cell cycle’ GO term differentially expressed (fold change>1.6x; 5%FDR) between HDF N and HDF 87y.

### FoxM1 repression dictates mitotic decline during natural aging

RNA-seq analysis also disclosed Forkhead box M1 (FOXM1), the transcription factor that primarily drives the expression of G2/M cell cycle genes, to be downregulated in elderly mitotic cells (logFC:-0.85; FDR<5%; Supplementary Table 1). Indeed, 66 out of the 105 ‘mitotic cell cycle’ genes altered in the octogenarian fibroblasts have been reported as targets of the Myb-MuvB (MMB)-FOXM1 transcription complex containing the cell cycle genes homology region (CHR) motif in their promoters^38,39^ (Supplementary Fig.6a). In addition, loss of FOXM1 has previously been shown to cause pleiotropic cell-cycle defects leading to embryonic lethality^40^. We therefore asked whether loss of mitotic proficiency during normative aging was due to FOXM1 downregulation. Indeed, increasing age correlated with decreased FoxM1 transcript and protein levels in our human (Fig. 4a-c) and mouse (Supplementary Fig.2f) samples. Importantly, unlike previous studies linking FoxM1 repression and aging 41,42, by analyzing the transcriptomes of mitotic cells only, we uncoupled FoxM1 downregulation from the subsequent reduction in cell proliferation, demonstrating that elderly cells divide with intrinsically low levels of FoxM1. We thus conclude that aging-associated FoxM1 repression likely accounts for the transcriptional shutdown of mitotic genes and the observed phenotypes in pre-senescent dividing cells.

**Figure 4.**
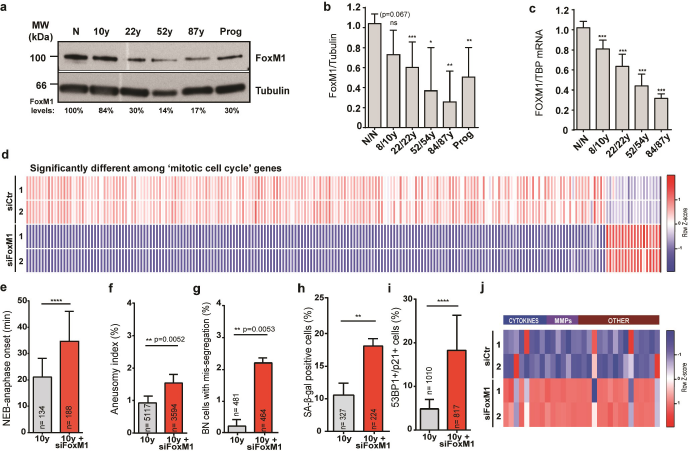
FoxM1 repression dictates cellular phenotypes associated with aging. **(a,b)** FoxM1 protein levels in mitotic extracts of different age samples. FoxM1 levels were normalized to -tubulin levels and to neonatal (N/N). **(c)** *FOXM1* transcript levels in mitotic total RNA from different age samples, normalized to *TBP* transcript levels and compared to neonatal. **(d)** Heatmap of genes within the ‘mitotic cell cycle’ GO term differentially expressed (fold change>1.6x; 5%FDR) by RNA-seq between control and FoxM1 siRNA-depleted 10 years-old fibroblasts. Genes are represented in columns and sample replicates are represented in rows. In the Z-score column, color intensities represent higher (red) to lower (blue) expression. **(e)** Mitotic duration (NEB-anaphase onset) of control and FoxM1 siRNA-depleted fibroblasts. **(f-g)** Aneusomy index in interphase cells **(f)** and in binucleated cells **(g)**. **(h-i)** Percentage of cells staining positive for the senescence markers β-galactosidase **(h)** and 53BP1/p21 **(i)**. **(j)** Heatmap of SASP genes differentially expressed between HDF 10y and 10y siFoxM1. Values are mean ± s.d. of three independent experiments. Sample size (n) is indicated in each graph. *p≤0.05, **p≤0.01, ***p≤0.001 and ****p≤0.0001 by Mann-Whitney (b,c,e) and two-tailed χ^2^ (f, g, h, i) statistical tests.

To test the correlation between FoxM1 repression and age-associated phenotypes further, we RNAi-depleted FoxM1 in fibroblasts from the 10 years-old donor (Supplementary Fig.5a,b) and performed RNA sequencing (RNA-seq) gene expression profiling of cells captured in mitosis. RNA-seq revealed that FoxM1 repression significantly altered the abundance of 4,451 gene transcripts (Supplementary Fig.6b,c; Supplementary Table 1), which were enriched for ‘mitotic cell cycle’ genes (206 genes, Gene Ontology [GO] analysis; 5% FDR) (Fig. 4d). This constituted a 2.52-fold enrichment of alterations in this gene set (*p*-value=1.09E-42). The top ten most altered GO terms included five related to mitosis (Supplementary Fig.6e), and 217 out of 249 G2/M cell cycle genes previously reported as targets of the DREAM (DP, RB-like, E2F4 and MuvB) and MMB-FOXM1 transcriptional complexes^43^, were downregulated following FoxM1 depletion (Supplementary Fig.6e). In agreement with these transcriptional alterations, FoxM1-depleted young fibroblasts displayed a mitotic delay (Fig. 4e), mitotic defects (Supplementary Fig.5c-e) and increased aneuploidy (Fig. 4f,g) similarly to old-aged fibroblasts. Interestingly, the percentage of cells with SA biomarkers (Fig. 4h,i), as well as the expression of SASP genes itself (Fig. 4j) also increased following FoxM1 repression. Altogether, these results show that young cells with low levels of FoxM1 recapitulate the aging-associated mitotic defects, aneuploidy and senescence.

### Constitutively active FoxM1 rescues mitotic decline and cellular senescence in elderly and HGPS cells

As FoxM1 depletion reduced mitotic fidelity and induced senescence in young cells, we hypothesized that FoxM1 overexpression in elderly cells should counteract aging phenotypes. Expression of a constitutively active truncated form of FoxM1 (FoxM1-dNdK) in old-aged fibroblasts^44,45^ (Supplementary Fig.5f,g) resulted in altered expression of 1,955 gene transcripts as determined by RNA-seq (Supplementary Fig.6b,d; Supplementary Table 1), including 85 ‘mitotic cell cycle’ genes (Fig. 5a). Moreover, 149 out of the 249 DREAM/MMB-FOXM1 gene targets were significantly upregulated following FoxM1-dNdK expression (Supplementary Fig.6e). In addition, we found extensive overlap between genes of mitosis-related GO terms that were downregulated in FoxM1 RNAi and upregulated in FoxM1-dNdK expression experiments (1,163 genes; Supplementary Fig.6e; Supplementary Table 2). Concordantly, the age-associated mitotic defects were ameliorated in elderly fibroblasts expressing FoxM1-dNdK (Fig. 5b,c; Supplementary Fig.5h-j), resulting in decreased aneuploidy levels (Fig. 5d,e). Furthermore, the percentage of cells with SA biomarkers (Fig. 5f,g) and the transcript levels of SASP genes (Fig. 5h) were decreased. Importantly, FoxM1-dNdK expression also partly rescued the mitotic defects, aneuploidy levels and percentage of senescent cells in HGPS fibroblasts (Fig. 5i-p). Overall, these data demonstrate that modulation of mitotic efficiency through FoxM1 induction in elderly and HGPS cells prevents both aneuploidy and cellular senescence.

**Figure 5.**
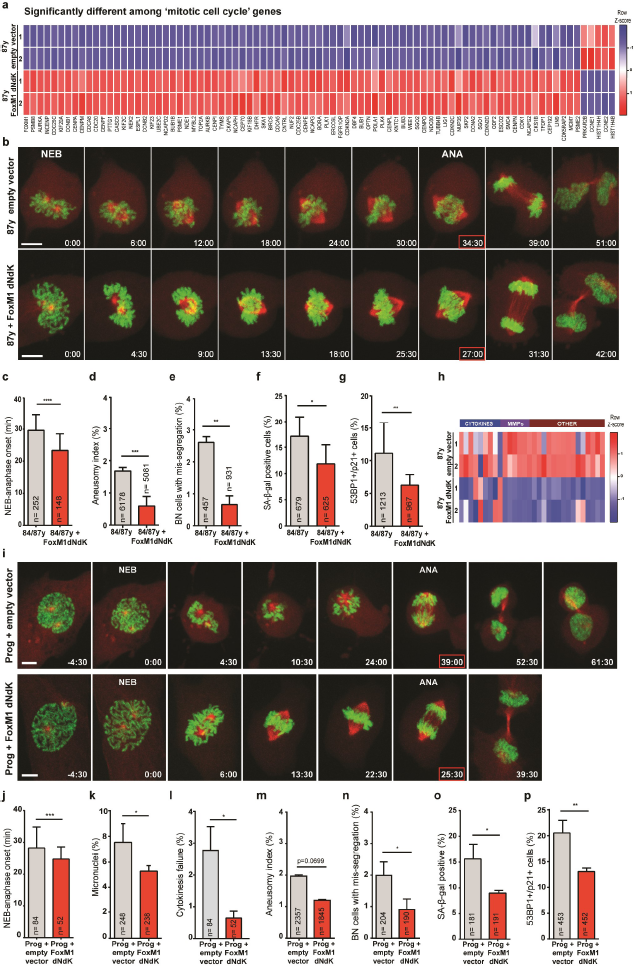
Constitutively active FoxM1 ameliorates mitotic fitness and aging markers in elderly and HGPS cells. (a) Heatmap of genes within the ‘mitotic cell cycle’ GO term differentially expressed (fold change>1.6x; 5%FDR) between mitotic elderly cells transduced with lentiviral empty vector or vector expressing FoxM1dNdK. In the Z-score column, color intensities represent higher (red) to lower (blue) expression. (b) Movie frames of elderly dividing cells expressing H2B-GFP--Tubulin-mCherry (upper panel) and H2B-GFP--Tubulin-mCherry + FoxM1dNdK (lower panel) (Supplementary Video 2). Time, min:sec. Scale bar, 5 µm. (c) Mitotic duration (NEB to anaphase onset) in elderly fibroblasts infected with control and FoxM1dNdK lentiviruses. (d-e) Aneusomy index in interphase (d) and binucleated cells (e). (f-g) Percentage of cells staining positive for the senescence markers β-galactosidase (f) and 53BPl/p2l (g). (h) Heatmap of SASP genes differentially expressed between HDF 87y and 87y FoxM1dNdK. (i) Movie frames of mitotic HGPS fibroblasts expressing H2B-GFP--Tubulin-mCherry (upper panel) and H2B-GFP--Tubulin-mCherry+FoxM1dNdK (lower panel) (Supplementary Video 3). Time min:sec. Scale bar, 5 µm. (j) Mitotic duration of HGPS fibroblasts transduced with empty and FoxM1dNdK lentiviruses. (k) Percentage of cells with micronucleus. (l) Percentage of cells failing cytokinesis. (m) Aneusomy index in interphase nuclei and (n) mis-segregation events in binucleated cells estimated from FISH analysis of 3 chromosome pairs. (o,p) Percentage of cells staining positive for the senescence markers (o) βgalactosidase and (p) 53BP1/p2l. Values are mean ± s.d. of three independent experiments. Sample size (n) is indicated in each graph. *p≤0.05, **p≤0.01, ***p≤0.001 and ****p≤0.0001 by Mann-Whitney (c,j) and two-tailed χ^2^ (d-g; k-p) statistical tests.

### Aneuploidy is a hallmark of elderly senescent cells

Experimental modulation of FoxM1 levels in the young (FoxM1 RNAi) and elderly fibroblasts (FoxM1-dNdK) underlined aneuploidy and senescence to be inherently linked. Previous studies have found mouse models of chromosomal instability to exhibit premature aging^11,46^, and cellular models of CIN to induce senescence^47^. Though informative, these studies suffer the drawbacks that CIN-inducing single gene editing leads to higher aneuploidy levels than those accumulating during normal aging, and is a non-physiological stress either present from early development onward (mouse models) or acutely induced (cellular models). Whether the mild aneuploidy levels associated with global transcriptional shutdown of mitotic genes account for cellular senescence in natural aging remains unknown. To address this question, we FACS-sorted senescent cells from neonatal or elderly fibroblast cultures using SA-β-gal as a senescence marker and measured the aneusomy index of three chromosome pairs (Fig. 6a-f). As expected, the percentage of SA-β-gal positive cells was considerably higher in elderly vs. neonatal early passage cell cultures (Fig. 6a,b,e), and consistent with our quantitative analysis using fluorescence microscopy (Supplementary Fig.1d). The aneusomy indexes of SA-β-gal positive cells were significantly higher than those of unsorted controls (Fig. 6f). Extrapolation of the aneusomy indexes to 23 chromosome pairs indicated aneuploidy levels of ~25% and ~70% in neonatal and elderly cell subpopulations, respectively. Thus, our data indicate that senescent cells from old donors are significantly more aneuploid than senescent cells from young donors, suggesting aneuploidy as a major hallmark of cellular senescence in natural aging. To investigate whether FoxM1 repression is responsible for this difference, we next FACS-sorted SA-β-gal positive cells from siFoxM1-depleted neonatal cell cultures and from octogenarian cell cultures expressing FoxM1-dNdK (Fig. 6c-e). We observed a 4.9x increase of senescent cells, and a comparable 5.3x increase in the aneusomy index, following FoxM1 repression in cells from neonatal donors. Conversely, a 2.2x decrease of senescent cells, and a 1.5x decrease in the aneusomy index was found following FoxM1-dNdK expression. These results demonstrate that FoxM1 levels modulate aneuploidy-driven senescence.

**Figure 6.**
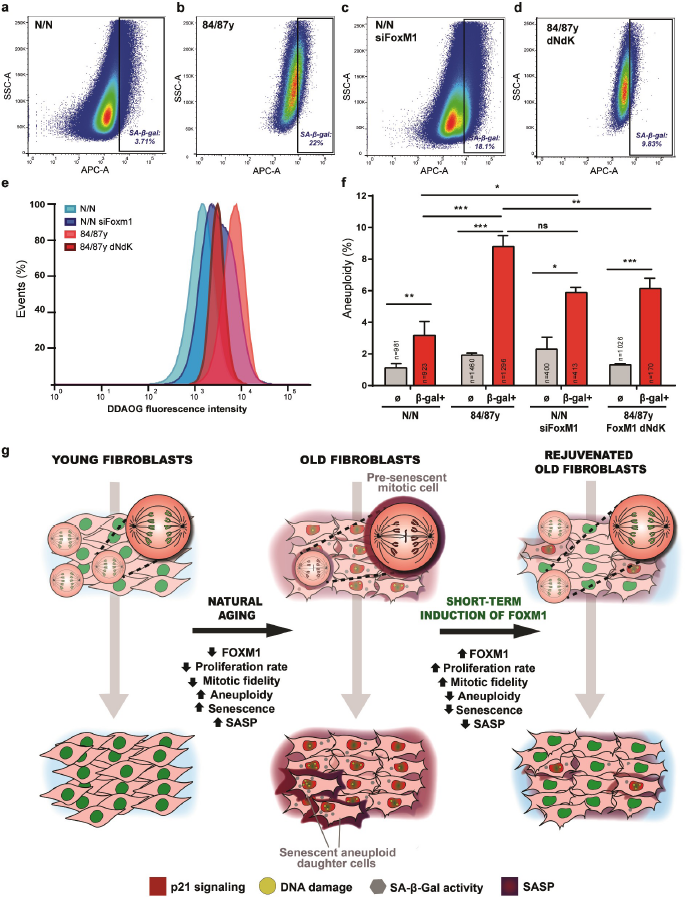
FoxM1 governs aneuploidization-driven cellular senescence in elderly cells. **(a-d)** FACS sorting of senescent cells from neonatal (a), elderly (b), FoxM1 siRNA-depleted neonatal (c) and 84y/87y with FoxM1dNdK (d) cell populations with high β-galactosidase activity. The gates were defined accordingly to the respective auto-fluorescent control. (e) Relative intensity levels of the fluorogenic substrate DDAOG in the sorted cell populations. (f) Aneuploidy index in FACS-sorted β-gal-positive fibroblast subpopulations (β-gal+) vs. unsorted populations () as determined by FISH analysis for 3 chromosome pairs. Values are mean ± s.d. of two independent experiments. Sample size (n) is indicated in each graph. ns: *p*>0.05, *p≤0.05, **p≤0.01, ***p≤0.001 and ****p≤0.0001 by two-tailed χ^2^ statistical test. (g) Model summarizing the molecular basis of age-associated mitotic decline and accumulation of senescent cells. FoxM1 repression in pre-senescent dividing cells leads to aneuploidy, which accounts for the transition into full senescence. FoxM1 induction in old cells restores mitotic fitness and erases aging markers.

Altogether, our data point up for a positive feedback loop between cellular aging and aneuploidy, in which naturally aged cells with early senescent phenotypes loose mitotic fidelity and generate an aneuploid progeny that primarily accounts for the accumulation of full senescent phenotypes. We show that induction of FoxM1 transcriptional activity in elderly cells ameliorates cell autonomous and non-autonomous feedback effects between mitotic fidelity and senescence (Fig. 6g).

## Discussion

Aging is the largest risk factor for most diseases^48^. In recent years, seminal studies have shown that the accumulation of senescent cells in tissues over time shortens healthy lifespan and that selective clearance of senescent cells can greatly postpone aging^32,49^. Our results bring novel insight into how senescent cells arise and provide a molecular basis to prevent senescence and thus, an alternative strategy to emergent senolytic therapies^50^.

*SASP phenotype in elderly mitotic cells and aneuploidy-driven senescence.* Previous studies have suggested G2 and 4N G1 (mitosis skip) as the phase at which senescent cells exit the cell cycle 51,52. However, these studies made use of the acute senescence stimuli, gamma irradiation and oncogenic Ras overexpression respectively. In contrast to acute senescence, chronic senescence results from long-term, gradual macromolecular damage caused by stresses during lifespan (e.g. loss of proteostasis, DNA damage, epigenetic changes). We demonstrate that in a natural aging cellular model, consisting of dermal fibroblasts retrieved from elderly donors and cultured under limited passage number to prevent replicative *in vitro* aging, senescent cells are mostly aneuploid as a result of defective chromosome mis-segregation. Nonetheless, as shown for oncogene-induced senescence and replicative senescence^52,53^, one round of cell division is likely required for a transition from ‘early’ to fully senescent state. Consistent with this idea of senescent state evolution, we found elderly mitotic cells to express SASP genes. The SASP is largely initiated by p38 MAPK signaling^54^, which has been shown to be activated by mitotic delay^55,56^, which fits with our observation that mitotic duration steadily increases with aging.

Even though aneuploidy has been correlated with senescence, previous studies have used genetically induced chromosomal instability models or *in vitro* models of acute senescence induction in which the aneuploidy levels are too high in comparison to the mild levels we found in our model of naturally aged cells. Nevertheless, using an innovative experimental setup, we demonstrated that FACS-sorted cells exhibiting high SA--gal activity are prominently aneuploid in old samples compared to young samples. We hypothesize this difference to be the result of an aneuploidization propensity of elderly cells, caused by the repression of players of the error-correction machinery for chromosome attachments (Supplementary Table 1).

*Molecular mechanism of aging mitotic decline.* We showed that the frequency of mitotic abnormalities increases in older cells resulting in mild aneuploidy levels. We identified FoxM1 transcription factor as the molecular determinant of this age-associated mitotic decline. FoxM1 repression is likely mediated by activation of stress pathways, presumably triggered by primary causes of cellular damage, such as genomic instability or loss of proteostasis. For instance, it has been shown that genotoxic stress can activate the p53-p21-DREAM pathway, which in turn prevents early cell cycle gene expression required for FoxM1 transcriptional activity^43,57^.

Transcriptional deregulation during aging has been widely reported in asynchronous cell populations^58,59^. In this study, we used synchronized mitotic populations and found that many cell cycle genes were deregulated in old cells when compared to young cells. Intriguingly, old mitotic cells have intrinsic low levels of mitotic transcripts. While this supports the observed mitotic abnormalities and increased aneuploidy, it is still surprising how elderly mitotic cells can cope with such reduced levels of transcripts (e.g. *CCNB1*). One possibility might be the balanced/stoichiometric repression of most mitotic genes. Furthermore, parallel mechanisms might buffer aging-mediated repression of mitotic transcripts. For example, even though we found SAC genes to be downregulated in elderly cells, SAC functionality was similar in young and old cells, suggesting that a parallel downregulation of genes contributing to proteasome activity (e.g. APC/C subunits) might buffer SAC gene repression (Supplementary Table 1).

*Modulation of FoxM1 activity to ameliorate cellular aging phenotypes.* We demonstrate that reestablishment of FoxM1 expression in elderly and HGPS fibroblasts can rescue mitotic decline and aneuploidy-driven senescence. Fibroblasts are the main constituents of connective tissue and are important players in extracellular matrix production and tissue repair. Therefore, preventing the accumulation of senescent fibroblasts will counteract a SASP-induced inflammatory microenvironment in these tissues, and will help to protect stem cell and parenchymal cell functions. Re-expression of FoxM1 could thus protect against adult stem cell and post-mitotic tissue aging. Indeed, increased expression of one FoxM1 transcriptional target, BubR1^43^, was previously shown to prevent aneuploidization and delay aging^13^. However, other aneuploidy mouse models have not been reported to exhibit premature aging. Possible explanations are the premature sacrifice of mice before they start developing aging phenotypes later in life and the cursory analysis for overt age-related degeneration missing tissue-specific phenotypes^14^.

Alternatively, aneuploidy-associated genes that are strongly linked with early aging might require induction of mild levels of aneuploidy or counteracting functions in additional cellular stresses that engage senescence response pathways. This appears to be the case of FoxM1. Not only FoxM1 repression translates into mild aneuploidy levels, but also FoxM1 might further act by counteracting age-associated cellular damage caused by genotoxic and oxidative stresses^42,60^.

Interestingly, increased FoxM1 expression was shown to improve liver-regenerating capacity in older mice^41^ and lung regeneration following injury^61^ without being tumorigenic, reinforcing that transcriptional modulation of mitotic fidelity could become a powerful therapy against aging and progeria syndromes, with advantages over other emergent strategies^50,62^. Not only it would circumvent the risks associated with cellular reprogramming to pluripotency, but it would also prevent senescent cell accumulation and thus avoid any irreversible deterioration of tissue extracellular matrix, which can not be rescued by senolysis^32^. Thus, our findings disclose a molecular mechanism with clinical benefit to healthy lifespan extension and HGPS treatment.

## Acknowledgments

All data are archived at the i3S Institute. We thank H. Maiato, J.M. Cabral and J. Bessa at i3S for critical discussions and reading of the manuscript; P. Sampaio and A. Maia for technical help with Advanced Light Microscopy. E.L. holds an FCT Investigator Postdoctoral Grant (IF/00916/2014) from FCT/MCTES (Fundação para a Ciência e a Tecnologia/Ministério da Ciência, Tecnologia e Ensino Superior). FCT Fellowship (SFRH/BD/74002/2010) supported J.C.M. The following project grants supported this work: PTDC/BEX-BCM/2090/2014 funded by FCT; NORTE-01-0145-FEDER-000029 funded by North Regional Operational Program (NORTE2020) under PORTUGAL 2020 Partnership Agreement through Regional Development Fund (FEDER); NORTE-07-0124-FEDER-000003 co-funded by North Regional Operational Program (ON.2) through FEDER and by FCT; and POCI-01-0145-FEDER-007274 i3S framework project co-funded by COMPETE 2020/PORTUGAL 2020 through FEDER and by FCT; Foundation Pediatric Oncology Groningen grant and Dutch Cancer Society grant 2012-RUG-5549 to FF.

## Author Contributions

J.C.M. initiated the project. J.C.M, S.V. and R.R. designed and performed the experiments, and analyzed the data. B.B. and P.B. performed RNA-sequencing and bioinformatics analysis. J.M.E. performed telomeric-FISH analysis. M.G.F., R.M. and F.F. contributed to the study design and edited the manuscript. E.L. conceived the idea, supervised the work and wrote the manuscript. All authors discussed results, prepared Fig.s and edited the manuscript.

## Competing Financial Interest

The authors declare no competing financial interests.

Correspondence and requests for materials should be addressed to E.L.

## Methods

### Cell culture

A total of 11 human fibroblast cultures, established from skin samples of Caucasian males with ages ranging from neonatal to octogenarian (two biological samples per age), were acquired from cell biobanks as summarized (Supplementary Fig.1A). Several time points over the human lifespan were included to reinforce the validity of any correlation found. All donors were reported as “healthy”, except the 8 years-old donor diagnosed with the Hutchison-Gilford progeria. Human dermal fibroblasts (HDFs) were seeded at 1 × 10^4^ cells per cm^2^ of growth area in minimal essential medium Eagle-Earle (MEM) supplemented with 15% fetal bovine serum (FBS), 2.5 mM L-glutamine and 1x antibiotic-antimycotic (all from Gibco, Thermo Fisher Scientific, CA, USA). Only early passage dividing fibroblasts (population doubling PDL<24) were used in all experiments. *PDL=3.32 (log UCY - log l) + X*, where UCY = the cell yield at that point, l = the cell number used to begin that subculture, and X = the doubling level of the cells used to initiate the subculture being quantitated. Murine adult fibroblasts (MAFs) were cultured in Dulbecco’s Modified Eagle Medium (DMEM) supplemented with nutrient mixture F-12, 10% FBS, L-glutamine and antibiotic-antimycotic (all from Gibco). All cells were grown at 37 °C and humidified atmosphere with 5% CO_2_.

### Isolation of mouse fibroblasts

Sv/129 mice were housed and handled accordingly to European Union and national legislation. Murine adult fibroblasts were collected from ears of n>3 sv/129 females of the same litter at their age of 8 weeks, 6 months, 1 year, 1.5 years and 2 years. Ears were washed with phosphate-buffered saline (PBS), cut into small pieces, and incubated with 1 mg/ml collagenase D and 1 mg/ml collagenase/dispase (both from Roche Applied Science, Germany) in DMEM:F12 without FBS, for 45 min at 37 °C and 5% CO_2_. Cells were then grown on a 6-well dish containing DMEM:F12, supplemented with 10% FBS and antibiotic-antimycotic.

### Drug treatments

Fibroblasts were incubated for 24 h in medium containing 2 µg/ml cytochalasin D (C8273, Sigma-Aldrich, MO, USA) to block cytokinesis. For SAC inhibition, cells were treated with 5 µM Mps1 inhibitor^63^ (AZ3146, TOCRIS, USA). For proteasome activity enhancement, 10 µM of Usp14 inhibitor^64^ were used (I-300, BostonBiochem, Cambridge, MA). To measure spindle mispositioning in metaphase cells, proteasome was inhibited with 5 µM MG132 (474790, Calbiochem, CA, USA) for 1.5 h. To inhibit kinesin-5, S-Trityl-L-cysteine (STLC) (2799-07-7, Tocris, USA) was used at different concentrations depending on the experiment: 5 µM during 16 h to enrich the mitotic index for mitotic cell shake-off, and 2.5 µM for 2 h if followed by a washout. Aurora kinase inhibitor ZM447439 (2458, AstraZeneca, Alderley Park, UK) was used at 500 nM for 18 h to artificially induce aneuploidy^65^.

### Fluorescence in situ hybridization (FISH)

Fibroblasts were grown on Superfrost™ Plus microscope slides (Menzel, Thermo Scientific, CA, USA) placed in a quadriperm dish (Sarsted, Nümbrecht, Germany). In both interphase and binucleated cell FISH, cells were given a hypotonic shock during 30 min (0.03 M sodium citrate, Sigma-Aldrich, MO, USA), followed by fixation in ice-cold Carnoy fixative added drop-wise and incubated for 5 min. This step was repeated two more times. FISH was performed with the Vysis centromeric probes CEP7 Spectrum Aqua, CEP12 Spectrum Green and CEP18 Spectrum Orange (Abbott Laboratories, Chicago, IL, USA) according to manufacturer’s instructions. Slides were mounted with mounting medium containing DAPI (Vectashield, Vector Laboratories, CA, USA).

### Senescence-associated β-galactosidase assay

In fixed cell analysis, cells were incubated for 90 min in medium with 100 nM Bafilomycin A1 (B1793, Sigma-Aldrich, MO, USA) to induce lysosomal alkalinization. 33 μM of fluorogenic substrate for β-galactosidase, fluorescein di-β-D-galactopyranoside (F2756, Sigma-Aldrich, MO, USA) were then added to the medium, and incubation carried out for 90 min. Cells were fixed in 4% paraformaldehyde for 15 min, rinsed with PBS and permeabilized with 0.1% Triton-X100 in PBS for 15 min. Finally, cells were counterstained with 1 µg/ml DAPI (Sigma-Aldrich, MO, USA). For fluorescence-activated cell sorting, cells were incubated in Bafilomycin A1 as described above and then exposed to 10 μM of fluorogenic substrate for β-galactosidase, 9H-(1,3-Dichloro-9,9-Dimethylacridin-2-One-7-yl) β-D-Galactopyranoside (DDAOG) for 90 min (Setareh Biotech LLC, USA).

### Fluorescence-activated cell sorting (FACS)

FACS was used to isolate subpopulations of senescent (SA-β-gal positive) live fibroblasts. FACS sorting was performed in FACSAria™ I Cell Sorter (BD Biosciences, CA, USA), using the laser-line of 633 nm. All cells within a single experiment were detected using the same voltage settings and sorted using an 85 μm nozzle. Cells were initially gated by forward scatter-area (FSC-A) vs. side scatter-area (SSC-A), which excludes dead cells and subcellular debris, with subsequent exclusion of cell doublets and clumps through FSC-A vs. FSC-width (FSC-W) plot. The signal was detected using the APC-A channel. The relative β-galactosidase activity was inferred from the median fluorescence intensity of the population. The sorting gates were designed accordingly to the respective auto-fluorescent control. Cells were sorted directly into MEM with 15% FBS, and seeded in Superfrost™ Plus microscope slides for subsequent FISH analysis. Analysis of FACS-data was done using FlowJo v10 software (TreeStar, Inc, Ashland, OR).

### Immunostaining

Fibroblasts were grown on sterilized glass coverslips coated with 50 µg/ml fibronectin (F1141, Sigma-Aldrich, MO, USA). For analysis of 53BP1 and Cdkn1a/p21 biomarkers, cells were fixed in freshly prepared 4% paraformaldehyde in PBS for 20 min, whereas for spindle angle analysis, cells were fixed in ice-cold methanol for 4 min. Following fixation, cells were rinsed in PBS and permeabilized in PBS + 0.3% Triton-X100 for 7 min. Cells were next blocked in 10% FBS in PBS-T (PBS + 0.05% Tween-20) for 1 h, and then incubated overnight at 4 °C with primary antibodies diluted in PBS-T + 5% FBS as follows: rabbit anti-53BP1 (#4937, Cell Signaling Technology, MA, USA), 1:100; mouse anti-p21 (SC-6246, Santa Cruz Biotechnology, CA, USA), 1:1,000; mouse anti-α-tubulin (T5168, Sigma-Aldrich, MO, USA), 1:1,500; rabbit anti-CP110 (a gift from Mónica Bettencourt-Dias, IGC, Portugal), 1:100. Secondary antibodies AlexaFluor-488 and -568 (Life Technologies CA, USA) were diluted 1:1,500 in PBS-T + 5% FBS. DNA was counterstained with 1 µg/ml DAPI (Sigma-Aldrich, MO, USA). Coverslips were mounted in slides with mounting solution (90% glycerol, 0.5% N-propyl-gallate, 20 nM Tris, pH 8).

### Telomere PNA FISH

Fibroblasts were incubated in 0.05 µg/ml colcemid (15212012, Gibco, Thermo Fisher Scientific, CA, USA) for 4 h, to induce metaphase arrest. Following trypsinization, fibroblasts were incubated in 0.03 M sodium citrate for 30 min at 37 ºC. Cells were then fixed in freshly made Carnoy fixative solution and stored at 4 °C. Metaphase spreads were fixed with 4% formaldehyde in PBS for 2 min, followed by pepsin digestion (1 mg/ml) for 10 min at 37 ºC. After a dehydration step with ethanol, dried slides were hybridized with Telomere-PNA probe (Applied Biosystems, CA, USA) (0.5 µg/ml) in 10 mM Tris pH 7.5, 70% formamide, 0.25% blocking reagent (Roche, Germany), 2 mM MgCl_2_, 700 µM citric acid, 7 mM Na_2_HPO_4_, for 3 min in a hot plate at 80 ºC and then for 2 h at 37 ºC in humidified chamber. Slides were washed in 70% formamide, 10 mM Tris, 0.1% BSA twice for 15 min, then washed 3 times in Tris-buffered saline (TBS) for 5 min, and finally mounted with mounting media containing DAPI (Vectashield, Vector Laboratories, CA, USA).

### Microscopy and image analysis

*Phase-contrast live cell imaging.* Fibroblasts were grown in glass-bottom 35mm µ-dishes (Ibidi GmbH, Germany), coated with 50 µg/ml fibronectin (F1141, Sigma-Aldrich, MO, USA). Images were acquired on a Zeiss Axiovert 200M inverted microscope (Carl Zeiss, Oberkochen, Germany) equipped with a CoolSnap camera (Photometrics Tucson, USA), XY motorized stage and NanoPiezo Z stage, under controlled temperature, atmosphere and humidity. Neighbor fields (20-25) were imaged every 2.5 min for 2–3 days, using a 20x 0.3NA A-Plan objective. Stitching of neighboring fields was done using the plugin “Stitch Grid” (Stephan Preibisch) from ImageJ/Fiji software.

*Spinning-disk confocal microscopy.* Four-dimensional data sets were collected with Andor Revolution XD spinning-disk confocal system (Andor Technology, Belfast, UK), equipped with an electron multiplying CCD iXonEM Camera and a Yokogawa CSU 22 unit based on an Olympus IX81 inverted microscope (Olympus, Southend-on-Sea, UK). Two laser lines at 488 and 561 nm were used for the excitation of GFP and mCherry and the system was driven by Andor IQ software. Z-stacks (0.8-1.0 μm) covering the entire volume of the mitotic cells were collected every 1.5 min with a PlanApo 60x 1.4NA objective. All images represent maximum-intensity projections of all z planes. ImageJ/Fiji software was used to edit the movies.

*Fluorescence microscopy.* Analysis of the SA-β-galactosidase fluorescence assay was carried out on a Zeiss AxioImager Z1 (Carl Zeiss, Oberkochen, Germany) equipped with an Axiocam MR and using an EC-Plan-Neofluor 40x 1.3NA objective. Cells displaying >5 fluorescent granules were considered positive for SA-β-gal activity. Calculation of spindle angle in relation to growth surface was done in images acquired as optimal distance z-stacks on a Zeiss AxioImager Z1 using a Plan-Apochromat 63x 1.4NA objective. AutoQuant X2 (Media Cybernetics, Rockville, USA) was used for image deconvolution and generation of different axes projections.

*Automated microscopy.* Image fields of 53BP1/p21 double immunostaining and FISH staining were acquired on IN Cell Analyzer 2000 (GE Healthcare, UK), equipped with a Photometrics CoolSNAP K4 camera and using a Nikon 20x 0.45NA Plan Fluor objective. Fluorescence intensity thresholds were set by eye and used consistently for samples within each experiment.

*Image analysis.* Fixed cell experiments (FISH, SA biomarkers and micronuclei) were blindly quantified using ImageJ/Fiji software. Mitotic duration, asynchronous daughter cell adherence and cytokinesis failure were quantified from phase-contrast movies. Quantifications in Fig. 2e and Fig. 2g were from spinning-disk confocal movies.

### Western blotting

Mitotic cell populations were collected by shaking off cell culture flasks enriched for mitotic index (MI) by a 16 h treatment with STLC. MI>95% was determined by visual scoring of cells with condensed chromosomes after Carnoy fixation, followed by with 1 µg/ml DAPI in PBS. Lysis buffer (150 nM NaCl, 10 nM Tris-HCl pH 7.4, 1 nM EDTA, 1 nM EGTA, 0.5% IGEPAL) with protease inhibitors was added to mitotic cell pellets, and lysates quantified for protein content by the Lowry Method (DC™ Protein Assay, BioRad, CA, USA). 20 μg of total extract were then loaded in SDS-PAGE gels and transferred onto nitrocellulose membranes for western blot analysis. Membranes were blocked during 1 h with TBS containing 5% low-fat milk. Primary antibodies were diluted in TBS containing 2% low-fat milk as follows: rabbit anti-FoxM1 (#13147, ProteinTech Group Inc, IL, USA), 1:1,000; mouse anti α tubulin (T5168, Sigma-Aldrich, CA, USA), 1:50,000; and mouse anti-GAPDH (#60004, ProteinTech Group Inc, IL, USA), 1:30,000. Goat anti-rabbit (SC-2004, Santa Cruz Biotechnology, CA, USA) and goat anti-mouse (SC-2005, Santa Cruz Biotechnology, CA, USA) HRP-conjugated secondary antibodies were diluted at 1:3,000 in TBS containing 2% low fat milk. Signal was detected using Clarity Western ECL Substrate reagent (Bio-Rad Laboratories, CA, USA) accordingly to manufacturer’s instructions. A GS-800 calibrated densitometer with Quantity One 1-D Analysis Softwar 4.6 (Bio-Rad Laboratories, CA, USA) was used for quantitative analysis of protein levels.

### Lentiviral plasmids

H2B-GFP^66^ was amplified as a B*gl*II-H2B-GFP-T2A-B*am*HI-N*ot*I fragment. This PCR fragment was digested with BglII+NotI and ligated into pRetrox-Tight-Puro (Clontech, CA, USA) digested with BamHI+NotI, thus destroying the 5’ BamHI/BglII site while reintroducing a BamHI site 3’ of H2B-GFP-T2A. In parallel, -Tubulin^66^ was amplified as a BglII--TubulinB*am*HI-N*ot*I fragment and ligated into pRetrox-Tight-Puro digested with B*am*HI+N*ot*I, again destroying the 5’ BamHI/BglII site while reintroducing a BamHI site 3’ of -Tubulin. mCherry was amplified from pExchange-1-Cherry (Agilent Technologies, CA, USA) as a B*gl*II-mCherryN*ot*I fragment and was ligated into pRetrox--Tubulin digested with BamHI-NotI, yielding pRetrox--Tubulin-mCherry. Finally, -Tubulin-mCherry was PCR-amplified as a BglII--Tubulin-mCherry-NotI fragment and ligated into pRetrox-H2B-GFP-T2A digested with BamHI+NotI, yielding pRetrox-H2B-GFP-T2A--Tubulin-mCherry. To obtain pLVX-Tight-Puro-H2B-GFP-T2A--tubulin-mCherry, a B*gl*II-H2B-GFP-T2A--Tubulin-mCherry-N*ot*I fragment was amplified from pRetroX-H2B-T2A-GFP--Tubulin-mCherry and ligated into pLVX-Tight-Puro (Clontech, CA, USA) digested with *BamHI+NotI*. To generate pLVX-Tight-Puro-FoxM1dNdK, a BglII-FOXM1-dNdK-NotI fragment was amplified from pcDNA3-Flag-ΔN-ΔKEN-FoxM1 44 and ligated into pLVX-Tight-Puro digested with *BamH*I+*Not*I. Primers used are described in Supplementary Table 3.

### Lentiviral production and infection

Lentiviruses were produced according to the protocol described in Lenti-X Tet-ON Advanced Inducible Expression System (Clontech). Lentiviruses carrying empty pLVX-Tight-Puro, pLVX-Tight-Puro-H2B-GFP--Tubulin-mCherry or pLVX-Tight-Puro-FoxM1dNdK, as well as lentiviruses carrying pLVX-Tet-On Advanced (which expresses rtTA), were generated in HEK293T helper cells transfected with packaging plasmids (pMd2.G and psPAX2) using Lipofectamine 2000 (Life Technologies, Thermo Scientific, CA, USA). Human fibroblasts were co-infected for 12-16 h with responsive and transactivator lentiviruses at 2:1 ratio, in the presence of 8 µg/ml polybrene (AL-118, Sigma-Aldrich, MO, USA). In the following day, 750 ng/ml doxycycline (D9891, Sigma-Aldrich, MO, USA) was added to the medium to induce co-transduction. Phenotypes were analyzed and quantified 48-72 h later, and transfection efficiency monitored by scoring the number of fluorescent cells or by western blotting.

### FoxM1 RNA interference

Cells were transfected 1 h after plating, with 45 nM FoxM1 siRNA (SASI_Hs01_00243977 from Sigma-Aldrich, MO, USA) using Lipofectamine RNAiMAX (Thermo Scientific, CA, USA) according to manufacturer’s instructions. Phenotypes were analyzed and quantified 72 h post transfection, and depletion efficiency monitored by western blotting.

### Real-time PCR

Total RNA was obtained from mitotic fibroblasts (collected as depicted in Fig. 3a and in the Western blotting section) using the RNeasy^®^ Mini kit (Qiagen, Hilden, Germany). RNA purity and integrity was confirmed in the Experion™ system (Bio-Rad Laboratories, CA, USA). cDNA was synthesized from total RNA (1 μg) using iScript Advanced Select cDNA Synthesis kit (BioRad Laboratories, CA, USA). The 2-∆∆Ct method was used to quantify the transcript levels of FOXM1 against the transcript levels of the housekeeping gene (TBP/GAPDH). Primers were designed to span at least one exon-intron junction (Supplementary Table 3). Amplification was performed in a C1000 Touch Thermal Cycler (CFX384 Real-Time System, Bio-Rad Laboratories, CA, USA), and analyzed using CFX Manager Software (Bio-Rad Laboratories, CA, USA).

### RNA sequencing and bioinformatics

RNA was isolated from mitotic human fibroblasts and validated as described above for PrimePCR assay. RNA-sequencing libraries were prepared using TruSeq Stranded Total RNA with Ribo-Zero Human/Mouse/Rat (RS-122-2201; Illumina, CA, USA) according to manufacturer’s protocol. Pooled libraries were sequenced on an Illumina HiSeq 2500 (single-end 50 bp). Reads were aligned to the human genome (hg19) using a splicing-aware aligner (StarAligner). Aligned reads were FPM normalized, excluding low abundance genes (mean FPM>1). Differential gene expression analysis was performed using the R-package edgeR (v3.14.0 available from Bioconductor at http://www.bioconductor.org/packages/release/bioc/html/edgeR.html). Significant differential gene expression of aged fibroblasts was defined as p-value<0.05 and of genetic manipulations as a 2-logFC cutoff value <-0.7 or >0.7, FDR<0.05 and p-value<0.05 (Supplementary Table 1). For generation of RNA-seq heatmaps, FPM normalized read counts were used. Hierarchical clustering was performed using the R library heatmap.2. RNA-seq data of HDF N vs 87y represent two biological replicates of each biological sample from a single experiment. The RNA-seq data from FoxM1 RNAi and dNdK viral infection represent two technical replicates of each biological sample. For ontological analysis, differentially expressed genes were analyzed using Database for Annotation, Visualization and Integrated Discovery (DAVID) Functional Annotation Tool v6.8^67^ (www.david-d.ncifcrf.gov). To calculate the significance of gene set enrichment, empirical p-values were generated using DAVID tool. Genes differentially expressed following FoxM1 RNAi (dataset: 4451 genes) were overlapped with those differentially expressed following FoxM1dNdK expression (dataset: 2,199 genes). Overlap analysis was also performed with a dataset of 249 genes reported as targets of both DREAM and MMB-FoxM1 complexes^43^ (Supplementary Table 2). Gene ontology (GO) term enrichment analysis was performed using DAVID Functional Annotation Tool.

### Statistical Analysis

Sample sizes and statistical tests for each experiment are indicated in the Fig. legends. *p*-values were obtained using GraphPad Prism version 6 (GraphPad, San Diego, CA, USA). Data were tested for parametric vs. non-parametric distribution using D’Agostino-Pearson omnibus normality test. Spearman rank correlation, Mann-Whitney, two-tailed χ^2^-square or one-way ANOVA for multiple comparisons tests were then applied accordingly. ns: p>0.05, *p≤0.05, **p≤0.01, ***p≤0.001 and ****p≤0.0001. Values are shown as mean ± s.d. or mean ± s.e.m. When required, box-and-whisker plot with median, IQR and minimum and maximum values was used.

## Supplementary information

**Supplementary Figure 1.**
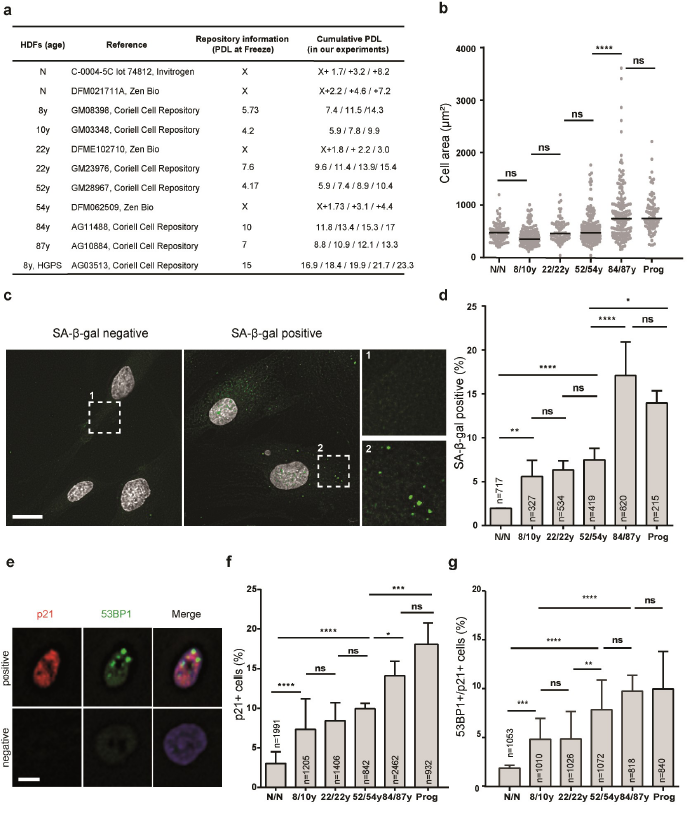
Senescence markers in human dermal fibroblasts from different age donors. **(a)** Fibroblasts from skin biopsies of Caucasian males with different ages used in this study. All donors were reported as “healthy”, except the 8 years old donor with the Hutchison-Gilford Progeria Syndrome. (b) Cellular area. Scatter plots show mean of n>50 cells. (c) Representative images of senescence-associated β-galactosidase (SA-β-gal) negative and positive cells. Scale bar, 20 µm. (d) Percentage of cells staining positive for senescence-associated β-galactosidase assay. (e) Representative images of cell cycle-arrested cells (p21 positive) and DNA damage (1 53BP1foci). Scale bar, 10 µm. (f) Percentage of cells staining positive for Cdkn1a/p21 cell cycle inhibitor. (g) Percentage of Cdkn1a/p21 positive cells with 53BP1foci. In all graphs, bars represent mean ± s.d. values from three independent experiments using two biological samples of similar age (except for progeria). Sample size (n) is indicated in each graph. ns: p>0.05, *p≤0.05, **p≤0.01, ***p≤0.001 and ****p≤0.0001 by two-tailed χ^2^ (d, f, g) and Mann-Whitney (b) statistical tests.

**Supplementary Figure 2.**
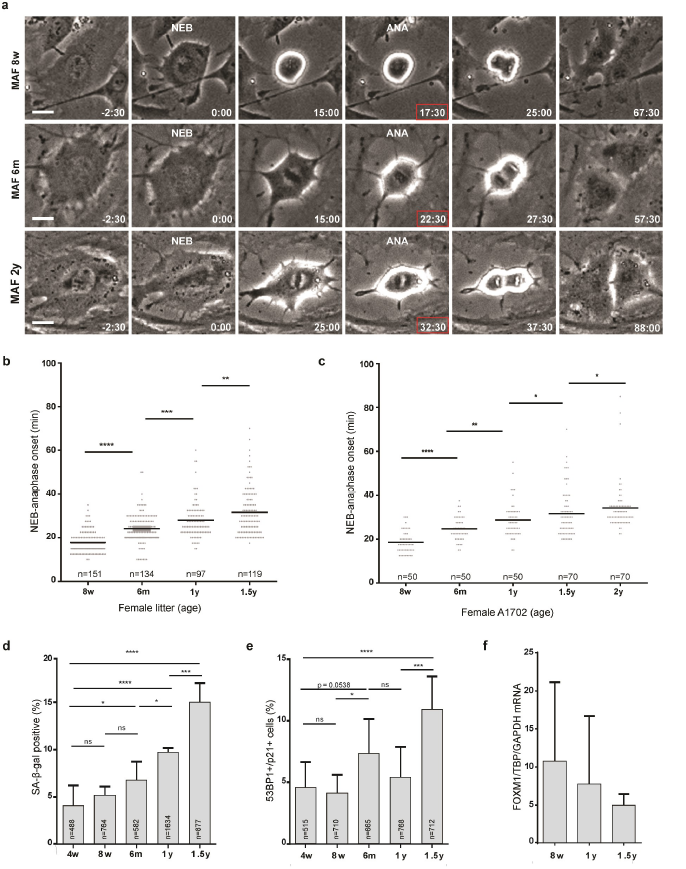
Aging-associated mitotic delay in mouse fibroblasts. **(a)** Movie frames of mitosis in MAFs collected at different age time-points as indicated. NEB, nuclear envelope breakdown. ANA, anaphase onset. Time min:sec. Scale bar, 20 µm. (b) Mitotic duration (NEB to anaphase onset) of individual MAFs collected at different age time-points from ≥3 sv/129 sister females. (c) Mitotic duration of individual MAFs collected at different age time-points from one randomly chosen single female (Al702). (d) Percentage of cells staining positive for senescence-associated β-galactosidase assay. (e) Percentage of Cdkn1a/p21 positive cells with 53BP1foci. **(f)** *FOXM1* transcript levels in mitotic total RNA from different age samples. Values are mean ± s.d. normalized to *TBP* and *GAPDH* transcript levels. Values are shown as mean, and sample size (n) is indicated in each graph. ns: p>0.05, *p≤0.05, **p≤0.01, ***p≤0.001 and ****p≤0.0001 by Mann-Whitney statistical test.

**Supplementary Figure 3.**
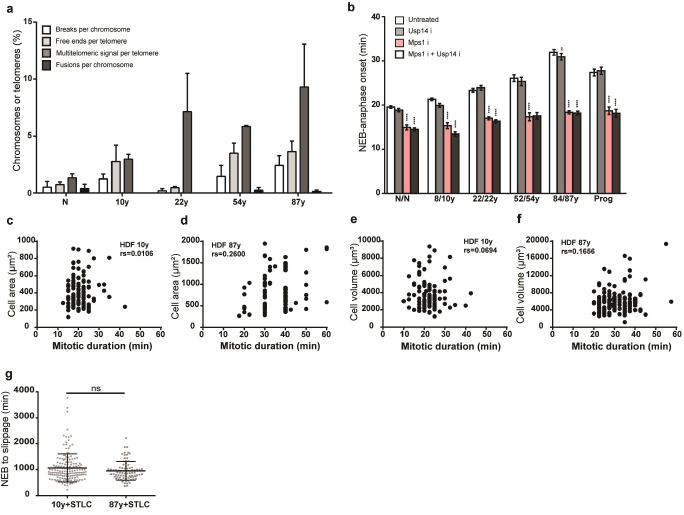
Effects of SAC activity, proteotoxic stress and cell size in mitotic duration. **(a)** Metaphase spreads of colcemid-treated cells of different age donors were quantified for chromosome fusions, chromosome breaks, telomere free ends and multitelomeric signals (MTS; marker of telomere replication defect). At least 900 chromosomes or 3,500 telomeres were quantified in two independent experiments. Values are shown as mean ± s.e.m. Age-dependent significant differences in chromosome aberrations were tested by one-way ANOVA for linear trend and multiple comparisons. Chromosome fusions, chromosome breaks and telomere free ends, not significant; MTS shown a linear trend with p-value≤0.05. (b) Mitotic duration of human fibroblasts from different age donors cultured under standard conditions (white bars) and following treatment with Usp14 inhibitor (light gray bars), Mps1 inhibitor (colored bars) and Mps1+Usp14 inhibitors (dark gray bars). (c-f) Spearman’s correlation coefficients (r_s_) between mitotic duration and (c-d) cell area or (e-f) cell volume. (g) Elapsed time from nuclear envelope breakdown (NEB) to cell’s adherence into the surface (slippage) in fibroblasts treated with kinesin-5 inhibitor (STLC). Values are mean ± s.d. of n>50 (b) and n>80 (c-g) scored cells from three independent experiments. ns: p>0.05, **p≤0.01, ***p≤0.001 by Mann-Whitney statistical test (b-g).

**Supplementary Figure 4.**
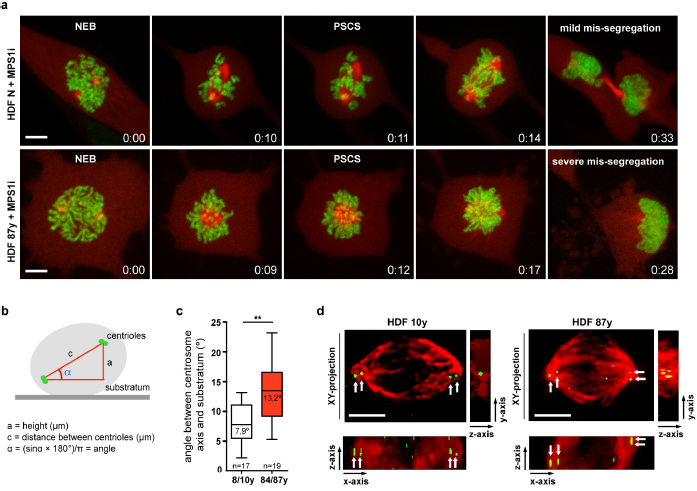
Quantitative analysis of aging-associated mitotic defects. **(a)** Chromosome alignment delay. Representative movie frames of neonatal and elderly cells inhibited for Mps1 to prematurely exit mitosis in the presence of misaligned chromosomes. Examples of mild chromosome mis-segregation (bipolar anaphase without cytokinesis failure) (upper panel) and severe chromosome mis-segregation (failure in anaphase movement and cytokinesis) (lower panel) are shown. NEB, nuclear envelope breakdown. PSCS, precocious sister chromatid separation. Time min:sec. (b-d) Spindle mispositioning. (b) Scheme depicting the calculation of the spindle angle in relation to the growth surface. (c) Box-and-whisker plot with median, IQR and minimum and maximum values, illustrating the range of spindle angle values from young and elderly metaphase cells. n indicates sample size. **p≤0.01 by Mann-Whitney test. (d) Orthogonal projections (in xy, xz and yz axes) of the 3D distribution of the spindle (α-tubulin, red) and centrioles (green) in young and elderly fibroblasts. Scale bar, 5 µm (a,d).

**Supplementary Figure 5.**
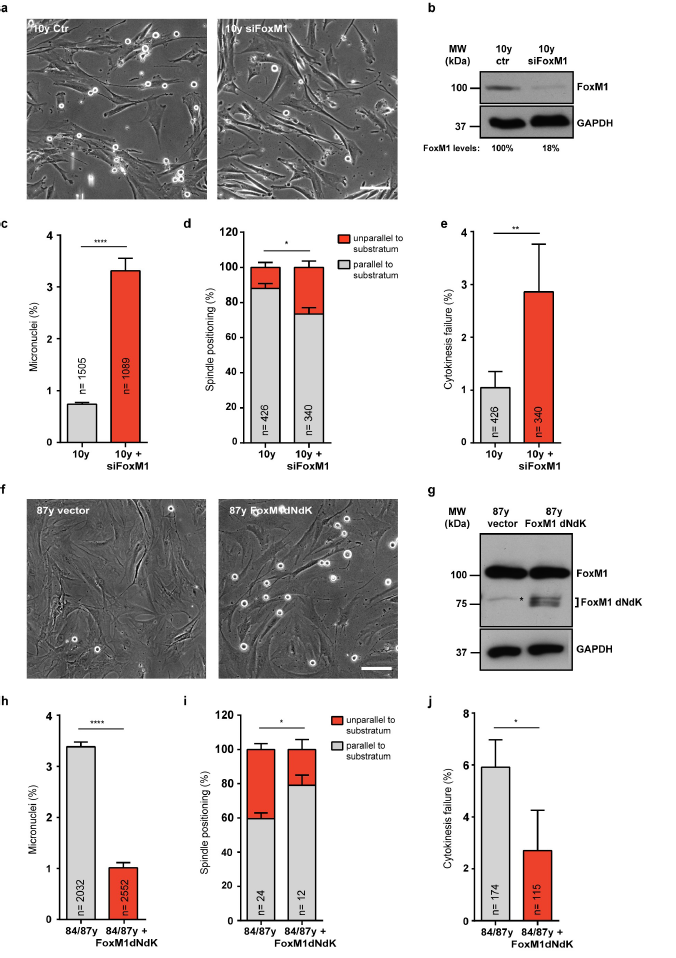
Modulation of FoxM1 levels affects mitotic fidelity. **(a-e)** FoxM1 repression in young cells. (a) FoxM1 RNAi depletion (siFoxM1) decreases mitotic index. (b) Western blot validation of FoxM1 decreased levels following RNAi. α-tubulin was used as loading control. (c) Percentage of cells with micronuclei. (d) Percentage of mitotic cells exhibiting unparalleled spindle to the substratum (in red). (e) Percentage of mitotic cells failing cytokinesis. (f-j) Expression of constitutively active FoxM1 (FoxM1dNdK) in elderly cells. (f) FoxM1dNdK expression increases mitotic index. (g) Western blot validation of FoxM1dNdK expression. Asterisk indicates unspecific band in untransduced 87y cells. GAPDH was used as loading control. (h) Percentage of cells with micronuclei. (i) Percentage of mitotic cells exhibiting unparalleled spindle to the substratum (in red). (j) Percentage of mitotic cells failing cytokinesis. Values are mean ± s.d. from three independent experiments. Sample size (n) is indicated in each graph. *p≤0.05, **p≤0.01 and ****p≤0.0001 by two tailed χ2 statistical test.

**Supplementary Figure 6.**
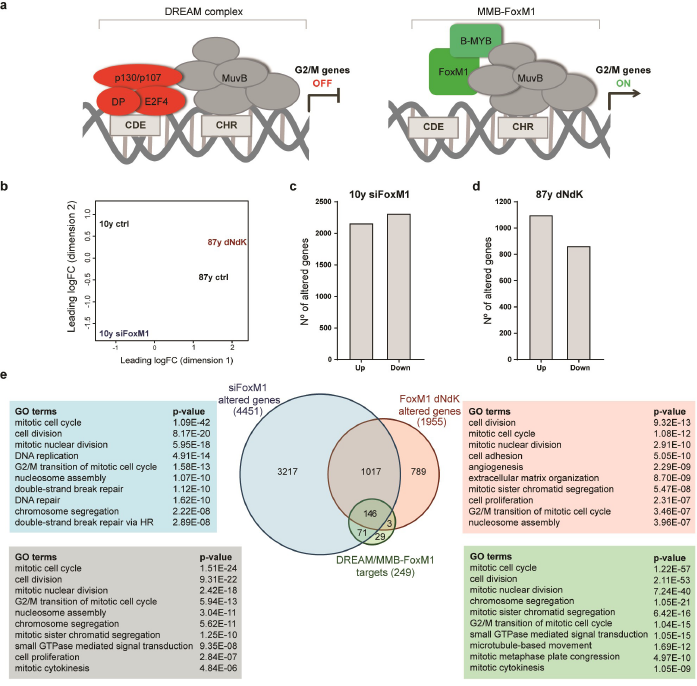
Meta-analysis of differentially expressed genes upon modulation of FoxM1 levels. **(a)** Regulation of G2/M cell cycle genes through sequential binding of DREAM and MMB-FoxM1 to CHR promoter elements, as described in^38^. (b) Multidimensional scaling plot of distances between gene expression profiles based on sample RNA-sequencing. (c) Number of altered genes upon FoxM1 RNAi in 10y cells. (d) Number of altered genes upon lentiviral transduction of 87y cells with FoxM1dNdK. Genes are listed in Supplementary Table 1. (e) Venn diagram displaying the overlaps between siFoxM1 altered genes, FoxM1dNdK altered genes and targets of both DREAM and MMB-FoxM1 complexes^43^ (Supplementary Table 2). The respective top ten GO terms are organized by p-values using the DAVID Functional Annotation tool.

Supplementary Table 1. RNA-sequencing data sets. Table includes comparative gene expression analysis.

Supplementary Table 2. Venn Diagram data sets. Overlap between FoxM1 RNAi altered genes, FoxM1dNdK altered genes, and DREAM/MMB-FoxM1 targets. Table includes all the overlapping clusters represented in the Venn diagram. Genes are listed accordingly to Ensembl ID.

Supplementary Table 3. Primers used in qPCR and cloning.

Supplementary Video 1. From left to right. Mitotic progression of a neonatal fibroblast (HDF N) co-expressing H2B-GFP (green) and α-tubulin-mCherry (red). Chromosome congression delay in a dividing elderly fibroblast (HDF 87y) co-expressing H2B-GFP (green) and α-tubulin-mCherry (red). Chromosome anaphase lagging and micronucleus formation in a dividing elderly fibroblast (HDF 87y) co-expressing H2B-GFP (green) and α-tubulin-mCherry (red). Unparalleled mitotic spindle positioning in relation to the growth surface and asynchronous adherence of the daughter cells in a dividing elderly fibroblast (HDF 87y) co-expressing H2BGFP (green) and α-tubulin-mCherry (red). Time-lapse images were acquired on a spinning-disk confocal microscope using 90 s intervals. Time min:sec. Frame series from this movie were used in Fig. 2a-d.

Supplementary Video 2. From left to right. Mitotic delay from nuclear envelope breakdown (NEB) to anaphase onset (ANA) in an elderly fibroblast (HDF 87y) co-expressing H2B-GFP (green) and α-tubulin-mCherry (red). Mitotic efficiency rescue following expression of FoxM1dNdK in an elderly fibroblast (HDF 87y) co-expressing H2B-GFP (green), α-tubulinmCherry (red). Time-lapse images were acquired on a spinning-disk confocal microscope using 90 s intervals. Time min:sec. Frame series from this movie were used in Fig. 5b.

Supplementary Video 3. From left to right. Chromosome alignment delay and mitotic spindle mispositioning in a HGPS (Progeria) fibroblast co-expressing H2B-GFP (green) and -tubulinmCherry (red). Mitotic efficiency rescue following FoxM1dNdK expression in a HGPS (Progeria) fibroblast co-expressing H2B-GFP (green) and -tubulin-mCherry (red). Time-lapse images were acquired on a spinning-disk confocal microscope using 90 s intervals. Time min:sec. Frame series from this movie were used in Fig. 5i.

